# Rapid phenotypic differentiation and local adaptation in Japanese knotweed s.l. (*Reynoutria japonica* and *R*. × *bohemica*, Polygonaceae) invading novel habitats

**DOI:** 10.1101/2022.03.07.483296

**Authors:** Wei Yuan, Massimo Pigliucci, Christina L. Richards

## Abstract

**PREMISE:** Many plant invaders like the Japanese knotweeds are thought to colonize new habitats with low genetic diversity. Such species provide an opportunity to study rapid adaptation to complex environmental conditions.

**METHODS:** Using replicate reciprocal transplants of clones across three habitats, we described patterns of phenotypic response and assessed degree of local adaptation.

**KEY RESULTS:** We found plants from beach habitats had decreased height, number of leaves, leaf area, and biomass allocation to roots and shoots compared to plants from marsh and roadside habitats when grown in their home habitat. In the marsh habitat, marsh plants were generally larger than beach plants, but not different from roadside plants. There were no differences among plants from different habitats grown in the roadside habitat. Despite this evidence of differentiation in beach and marsh habitats, we found mixed evidence for local adaptation. In their “home site” plants from the marsh habitat had greater biomass than plants from the beaches but not compared to plants from roadsides. Biomass comparisons in other habitats were either maladaptive or not significant. However, plants from the roadside had greater survival in their “home site” compared to foreign plants. There were no differences in survival in the other habitats.

**CONCLUSIONS:** We found phenotypic differentiation associated with habitats despite the low reported genetic diversity for these populations. Our results partially support the hypothesis of local adaptation in marsh and roadside habitats. Identifying whether these patterns of differentiation result from genetic or heritable non-genetic mechanisms will require further work.

## INTRODUCTION

The expansion of invasive species challenges our understanding of the process of adaptation given the likelihood of reduced genetic variation following a population bottleneck (Bock et al., 2015; Colautti and Lau, 2015). The classic population genetic assumption is that dramatically reduced genetic variation will severely constrain the evolutionary potential of a given population or species (Sakai et al., 2001; Allendorf and Lundquist, 2003). The chances of a few individuals or only one genotype landing in a novel location and surviving are expected to be minimal (Dlugosch and Parker, 2007; Bossdorf et al., 2008). Some invasive species benefit from alternative sources of increased genetic variation through multiple introductions (Durka et al., 2005; Lavergne and Molofsky, 2007; Rosenthal et al., 2008; Gammon and Kesseli, 2010; Qiao et al., 2019; Liu et al., 2020) or hybridization (Daehler and Strong, 1997; Bímová et al., 2001; Pysek et al., 2003; Mandák et al., 2004; Salmon et al., 2005; Bailey et al., 2009; Parepa et al., 2014; Qiao et al., 2019). However, many invasive species appear to do well even with low levels of sequence based variation (Hollingsworth and Bailey, 2000; Geng et al., 2007; Dlugosch and Parker, 2008; Loomis and Fishman, 2009). This could be perhaps through chance establishment of “general purpose genotypes” (Baker, 1965; Richards et al., 2006; Oplaat and Verhoeven, 2015), high performance genotypes, or rapid evolution of niche breadth (Matesanz and Sultan, 2013; Matesanz et al., 2015). Vankleunen *et al.* (van Kleunen et al., 2018) argue that adaptive evolutionary processes are at least as common in invasive species as native species, which is surprising given the short amount of time they have to evolve. In fact a meta-analysis of 134 plant species in 52 plant families showed that invasive plant species demonstrated patterns of local adaptation just as frequently, and at least as strongly as native plant species (Oduor et al., 2016).

Considering the limited genetic diversity of many invasive populations, phenotypic plasticity has often been suggested as a potentially important mechanism in the invasion process (Baker, 1965; Sexton et al., 2002; Sultan, 2004; Bossdorf et al., 2005). Several studies have found support for the importance of plasticity (Cheplick, 2006; Pan et al., 2006; Geng et al., 2007; Muth and Pigliucci, 2007; Bossdorf et al., 2008; Funk, 2008; Richards et al., 2008; Loomis and Fishman, 2009; Walls, 2010; Matesanz and Sultan, 2013; Sultan et al., 2013; VanWallendael et al., 2018), but see (Davidson et al., 2011). Further, epigenetic mechanisms may contribute to the persistence of plastic responses across generations (Herman and Sultan, 2016; Shi et al., 2018; Puy, Carmona, et al., 2021; Puy, de Bello, et al., 2021). Epigenetic effects have been shown to be especially important in response to hybridization and exposure to stressful or novel environments, which are circumstances often experienced by invasive plants (Mounger et al., 2021). The chance sampling of genotypes involved in the invasion process, combined with non-genetic sources of phenotypic variation, can lead to divergence in phenotypes of these populations even in the absence of abundant genetic variation (Keller and Taylor, 2008; Prentis et al., 2008; Neinavaie et al., 2021). Considering the plurality of potential mechanisms of rapid evolution in novel conditions, studies that examine phenotypic response of clonal replicates in natural settings will enhance our understanding of the processes of adaptation.

Invasive populations of the *Reynoutria* species complex (referred to as Japanese knotweed *sensu lato* or *s.l.)* are known to occupy a wide range of habitats in Europe (Pyšek et al., 2001; Mandák et al., 2004; Bailey et al., 2009; Zhang et al., 2016), and the United States (Barney, 2006; Grimsby et al., 2007; Richards et al., 2008, 2012; Walls, 2010; Gaskin et al., 2014). On Long Island, NY, in addition to roadside habitats and along railways, they are often growing on beaches and next to or on the terrestrial border of marshes that are inundated with *Phragmites australis* (Richards et al., 2008, 2012; Walls, 2010). These populations provide a unique opportunity to explore rapid adaptation to very different habitat types. The physiological challenges of living in beach and salt marsh habitats might require a dramatic phenotypic shift. For example, plants that are able to tolerate highly saline environments are characterized by traits that specifically ameliorate the toxic and osmotic effects of substrate salinity to allow for growth under these conditions (Flowers et al., 1977; Donovan et al., 1996; Rosenthal et al., 2002; Lexer et al., 2003; Karrenberg et al., 2006; Richards et al., 2010). We used cytology and AFLP markers to show that these populations have extremely low genetic diversity: some are made up of a single *R. japonica* genotype reported across the US and Europe (Hollingsworth and Bailey, 2000; Mandák et al., 2005; Grimsby and Kesseli, 2010; Krebs et al., 2010; Gaskin et al., 2014; Groeneveld et al., 2014; Zhang et al., 2016), while the majority consist of a few *R. × bohemica* hybrids (Richards et al., 2008, 2012; Walls, 2010). In the greenhouse, plants from both roadside and marsh habitats were highly plastic in response to treatment with salt, but within and among sites there were significant differences in most traits and trait plasticities (Richards et al., 2008).

While controlled experiments allow us to examine response to specific environmental factors, reciprocal transplantation in natural field settings is the most robust approach to identify local adaptation to complex real environments (Kawecki and Ebert, 2004). Plants are considered to be locally adapted if they perform better in their “local site” or “home site” than do plants from other “foreign” sites. A robust quantitative genetics design in a transplant study also allows for investigating which traits are under selection (Lande and Arnold, 1983; Dudley, 1996a; b; Mauricio and Rausher, 1997; Donovan et al., 2007), and characterizing the patterns of divergence of traits across different habitats (Roff and Mousseau, 2005; Doroszuk et al., 2008; Franks and Weis, 2008).

In this study, we used reciprocal transplants of Japanese knotweed to investigate 1) how traits vary in response to the different habitat types, and 2) if plants demonstrate patterns of adaptation to the different habitat types. If invasive Japanese knotweed plants have heritable variation for traits that increase fitness in these habitats, selection should act on these traits and lead to locally adapted populations. If these populations are adapted, we expect that they will outperform plants from other habitats, through greater biomass or greater survival, when grown in their local habitat (Kawecki and Ebert, 2004). To test this hypothesis, we established reciprocal transplants among beach, marsh and roadside habitats, planting replicates of each individual back into its home site and into the other two habitat types. We then asked two questions: Are there persistent differences in ecologically important traits in plants from different habitats? Do these populations display evidence of local adaptation?

## MATERIALS AND METHODS

### Reynoutria species complex –

Historically, the taxonomy of Japanese knotweeds has been complicated (for a review see Bailey and Conolly, 2000; Bailey and Wisskirchen, 2004; Schuster, Reveal, et al., 2011; Schuster, Wilson, et al., 2011; specifically for the introduction to New York State see Townsend, 1997; Del Tredici, 2017). Briefly, the two species *Reynoutria japonica* and *R. sachalinensis* were originally from Japan and were introduced to Europe in the mid 1800’s and the United States by the end of the 19th century (Del Tredici, 2017). Although *R. sachalinensis* (2n=44, 66, 88 and 132; Tiébré et al., 2007; Park et al., 2018) is distinct morphologically from *R. japonica* (2n=44 or 88), the two species are not differentiated in chloroplast DNA (cpDNA, Inamura et al., 2000). Hybridization between them appears to be rare in Japan, where they are not usually sympatric (but see (Park et al., 2018) for evidence of introgression). However, the hybrid *R. × bohemica* is common in the invasive range in Europe (Pysek et al., 2003; Bailey and Wisskirchen, 2004; Mandák et al., 2004; Bailey, 2013; Groeneveld et al., 2014; Parepa et al., 2014) and the U.S. (Forman and Kesseli, 2003; Gammon et al., 2007; Richards et al., 2008, 2012). In the U.S., studies in New England suggested that spread of all three taxa takes place through both vegetative and sexual reproduction (Forman and Kesseli, 2003; Gammon et al., 2007; Grimsby et al., 2007; Gammon and Kesseli, 2010). However, other studies in the U.S. on *R. japonica* report only the same single female genotype that has also been found throughout Europe (Richards et al., 2012; Gaskin et al., 2014; Groeneveld et al., 2014; but see VanWallendael et al., 2021).

### Collection sites and experimental gardens –

In mid-May 2005, we collected Japanese knotweed *s.l.* rhizomes for reciprocal transplant studies between beach, marsh and roadside sites across Suffolk County, Long Island, New York (Table 1; four sites of each habitat type for 12 total sites were used as sources for plant material and as locations of transplants). We created four groups of plants for reciprocal transplant in order to maximize the ability to replicate each genet in one site of each of the habitat types while still testing for superior performance in the original “local” site.

**Table 1.** Locations of origin sites for the three habitat types within each transplant garden group for the 12 sites (used as sources for plant material and as locations of transplants). The table provides the number of rhizomes from each site (with range of replicates) and total number of replicates per site, as well as the total number of rhizomes and replicates within the four reciprocal transplants. The table also indicates the total number of surviving plants from each origin site (and number of survivors in beach, marsh and roadside transplant gardens).

| Transplant garden group | Origin site | Source habitat type | City | Latitude | Longitude | No. of rhizomes (reps/rhizome) | Total no. ramets | Survivors |
| --- | --- | --- | --- | --- | --- | --- | --- | --- |
| 1 | MSH | Beach | Port Jefferson | 40 57.7 | 73 02.6 | 6 (9-21) | 81 | 35 (3,25,7) |
| 1 | CBH | Marsh | Port Jefferson | 40 57.2 | 73 02.7 | 5 (6-21) | 117 | 32 (2,24,6) |
| 1 | ST | Roadside | Smithtown | 40 51.5 | 73 12.6 | 7 (6-15) | 117 | 43 (0,27,16) |
| **Total for transplant 1** |  |  |  |  |  | **18** | **315** | **110 (35%)** |
| 2 | PJB | Beach | Port Jefferson | 40 57.9 | 73 03.2 | 7 (9-21) | 93 | 19 (0,5,14) |
| 2 | CMM | Marsh | Center Moriches | 40 48.0 | 72 46.4 | 8 (9-24) | 75 | 8 (0,0,8) |
| 2 | CMR | Roadside | Center Moriches | 40 48.0 | 72 46.4 | 8 (6-21) | 72 | 8 (0,0,8) |
| **Total for transplant 2** |  |  |  |  |  | **23** | **240** | **35 (15%)** |
| 3 | RPB | Beach | Rocky Point | 40 58.0 | 72 57.3 | 6 (6-21) | 147 | 37 (2,12,23) |
| 3 | WH | Marsh | Brookhaven | 40 46.2 | 72 53.9 | 8 (6-21) | 117 | 8 (0,4,4) |
| 3 | HL | Roadside | Rocky Point | 40 57.8 | 72 57.3 | 8 (6-21) | 111 | 35 (2,17,16) |
| **Total for transplant 3** |  |  |  |  |  | **22** | **375** | **80 (21%)** |
| 4 | HP | Beach | Southold | 41 05.2 | 72 26.7 | 8 (12-24) | 120 | 39 (11,6,22) |
| 4 | RHBH | Marsh | Riverhead | 40 54.2 | 72 37.1 | 8 (6-19) | 132 | 76 (22,29,32) |
| 4 | RHC | Roadside | Riverhead | 40 54.5 | 72 37.5 | 8 (9-21) | 105 | 50 (9,16,25) |
| **Total for transplant 4** |  |  |  |  |  | **24** | **357** | **165 (46%)** |

Due to the natural topography of Long Island, the beach sites are all located on the northern shore of Long Island, while the salt marsh sites and roadside sites are more evenly distributed around Suffolk County. The beach sites are separated by 1-65 km, the marsh sites are separated by 14–40 km, and the roadside sites are separated by 20–32 km. At each site, we collected approximately one meter of rhizome from seven (Beach 1) or eight genets (all of the other 11 sites) that were approximately 10 m apart, to maximize the chances of sampling different genotypes and to represent plant distribution at each site. We refer to each of these rhizomes as a separate “genet” because replicates cut from the same rhizome should have the same genotype. However, our previous studies show that most “genets” within a site also have the same AFLP haplotype and most likely belong to the same individual. Rhizomes were brought to the Stony Brook University greenhouse and cut into pieces of 4-8 grams fresh-weight (12 sites x 7 or 8 rhizomes per site for a total of 95 genets x 18-25 replicates = 2225 rhizome pieces). We planted the rhizome pieces in individual wells in 24-well flats with Pro-mix potting medium (Pro-mix Bx, Quakertown, Pennsylvania, USA), and approximately one teaspoon of slow release fertilizer (15-9-12 Osmocote Plus 8-9 month, Marysville, Ohio). The flats were placed in a temperature-controlled greenhouse under conditions approximating mid-summer in Suffolk County, Long Island and watered as needed to keep the soil moist. Day temperature was maintained at 30°C and night temperature at 25°C. We grew the plants in the greenhouse for approximately six weeks to allow for shoots to emerge from the rhizomes and grow to a height of approximately 10-15 cm.

The 12 sites were organized into four reciprocal transplant groups, each with one beach, one marsh and one roadside location. Based on the number of plants that emerged, we assigned an equal number of replicate pieces of each rhizome collected at each source site to be transplanted into its home site and into one site of each of the other two habitat types. Therefore, three to eight replicates of each rhizome were assigned randomly across five blocks for each of the three transplant habitats (4 replicate studies x 3 transplant habitats x 3 source habitats x 5-8 genets x 2-8 replicates = 1287 plants; Table 1). We prepared the transplant gardens by removing only above ground vegetation (typically other knotweed plants) to a height of less than 2 cm with a machete. The cleared area included a border of 15-30 cm outside of the transplant blocks.

Between June 16-21, we transplanted the five blocks for each transplant garden into the field (between 80-125 ramets per transplant garden). Plants were left to grow in the field from the summer of 2005 through the fall of 2006. We measured final height and final number of leaves on all plants and harvested above ground and below ground for all plants between 11-18 September 2006. Roots were harvested by carefully digging to unearth the entire root system. Roots were shaken to remove loose dirt in the field and thoroughly washed.

### Traits measured –

We measured traits related to salt tolerance and overall performance for each plant: height, total number of leaves, total leaf area (Li-Cor Model LI-3100 Leaf area meter: Li-Cor, Inc., Lincoln, Nebraska, USA), succulence (g water in all leaves/ cm^2^ total leaf area), shoot dry biomass, root dry biomass, root:shoot ratio based on dry biomass, and total biomass at final harvest. For each plant, all live leaf tissue at final harvest was used for calculating total leaf area and succulence. Plants were dried in a forced air oven at 60° C for at least 72 h to determine shoot, root and total dry biomass. We evaluated survival and biomass (as proxies for fitness) to assess the degree of adaptation. These taxa have extensive clonal growth and many individuals may not flower at all in the field, but persist and spread from year to year so biomass is an important indicator of fitness (de Kroon and Groenendael, 1997).

### Data Analysis –

We performed all statistical analyses in the R statistical programming environment version 4.0.0 (R Core Team, 2020), using the Linear-Mixed-Model (LMM) or Generalized Linear-Mixed-Model (GLMM) framework (lme4 package, Bates et al., 2015) and the Bayesian simulation package arm (Gelman and Su, 2020). We checked the residuals to assess the appropriateness of the model and performed data transformations on traits as appropriate: we did not transform succulence and final height, but we performed log 10-transformation on leaf area, and log 2 transformation on shoot, root and total biomass. For these traits we fitted LMM with the model: trait<lmer(trait~SOURCE.type+GARDEN.type+SOURCE.type:GARDEN.type+(1|Origin.site)+ (1|Transplant.site)+(1|genetfactor),data=data,REML=F).

We used GLMM to model the number of leaves, with the negative binomial distribution with the model: modlfnum<glmer.nb(lf.number~SOURCE.type+GARDEN.type+SOURCE.type:GARDEN.type + (1|Origin.site)+(1|Transplant.site) + (1|genetfactor),data=data, REML=F).

We did not model root to shoot ratio directly, but instead we used the ratio of the estimates of mean and variance for root and shoot to assess significance within the Bayesian framework (Korner-Nievergelt et al., 2015).

In the LMM and GLMM models, “SOURCE.type” is the origin habitat type (beach, marsh, road), “GARDEN.type” is the transplant habitat type (beach, marsh, road). These effects as well as their interaction terms were modeled as fixed effects. The origin site, the transplant site, and the individual genets (“genetfactor”) were initially included as random terms. To avoid overfitting, we removed random effect terms that effectively explained no variance. This was true for the genet term for all traits and for the “Origin.site” term, which was removed for number of leaves, succulence and shoot biomass (see Table S1 for final models).

We examined the correlation matrix for each model to evaluate auto correlation between terms. To test the significance of the fixed effects of “SOURCE.type” and “GARDEN.type”, we used 95% credible intervals (CrI), a Bayesian analogue of confidence intervals (Bolker et al., 2009). For each response variable, we obtained the model estimates from the back-transformed effect sizes. We calculated the associated 95% Credible intervals (CrI) for the modeled effects by performing 10,000 iterations of Bayesian simulation of the mean and variance of each estimate, using the sim function in the R package arm with non-informative priors (Korner-Nievergelt et al., 2015). For each model, we examined distributions of simulated fitted values predicted by the fixed effect terms. We used the values corresponding to the 2.5 and 97.5 quantiles of the distribution to designate the lower- and upper-boundary of the 95% CrI. If the CrI of a group did not overlap with the mean of the other group within transplant gardens, we considered the difference between these groups to be significant (sensu Bucharova et al., 2016, 2017).

In order to understand how much of the variance was explained by the random and fixed effects in our model, we used several approaches. First, we used the package r2_nakagawa: Nakagawa’s R2 for mixed models (Nakagawa and Schielzeth, 2013; Nakagawa et al., 2017) to determine the conditional R^2^ (the variance explained by both the fixed and random effects) and marginal R^2^ (the variance explained by the combined fixed effects). The random effect variances calculated in this package are the mean random effect variances, and appropriate for mixed models with nested random effects (Johnson, 2014). We also used the package rptR (Stoffel et al., 2017) to further evaluate the components of variance for each of the random effects separately (i.e., “Origin site” and “Transplant site”. We used the package commonalityCoefficients (Nimon et al., 2008) to examine the amount of variance explained by the separate fixed effects of the habitats of the source and transplant gardens (i.e., “SOURCE.type” and “GARDEN.type”). This approach does not include information about the random effects of origin site and transplant site which are nested within the source type and garden type. However, this approach is valuable for evaluating the relative contribution of each separate fixed effect.

Our design is constrained by the fact that origin site and transplant sites are nested within levels of “Transplant group” (see discussion here Long, 2021). To examine the importance of this design constraint, we also reran the LMER and GLMER models for each trait with the fixed term “Transplant group”. To properly test for the effects in this nesting design, we should ideally fit random intercepts for the sites nested within groups, but we did not have enough replication within groups to do so. We assume that fitting the fixed effect of the “Transplant group” also controls for the non-independence of the origin site and transplant site within groups (Long, 2021). By comparing the modeling with and without the fixed term of Transplant group, we evaluated how these random terms impact the main effects of interest which are the fixed effects of the habitats of the source and transplant gardens (i.e., “SOURCE.type” and “GARDEN.type”).

We tested for local adaptation with two fitness proxies: total biomass and survival. We ran a “local vs. foreign” test using the Bayesian fitted values for biomass obtained from the same LMM model: totalbiomass<-lmer(trait~SOURCE.type+GARDEN.type+SOURCE.type:GARDEN.type+ (1|Origin.site)+(1|Transplant.site)+(1|genetfactor),data=data,REML=F).

For each garden type we performed random pairwise contrasts as the differences between Bayesian fitted values of the local plants with those from foreign habitats. For survival, we performed random pairwise contrast by calculating the log of odds ratio between local and foreign plants. We reported the mean, 95% CrI, and percentage of contrasts showing superior performance of plants grown in their home site compared to plants from each of the other habitats as magnitude of local adaptation.

## RESULTS

### Phenotypic response to reciprocal transplants –

We found that analyses of all traits resulted in large credible intervals (CrI) around the estimates of the means within source-by-garden combinations (Figure 1). This could be due to the large variance among transplant sites (Table 2, Table S2). Despite this large variance, we found differences in responses depended on the source habitat and the garden habitat. For every trait, at least one comparison met our significance criteria based on nonoverlapping of CrI of a group with the mean of another group.

**Figure 1.**
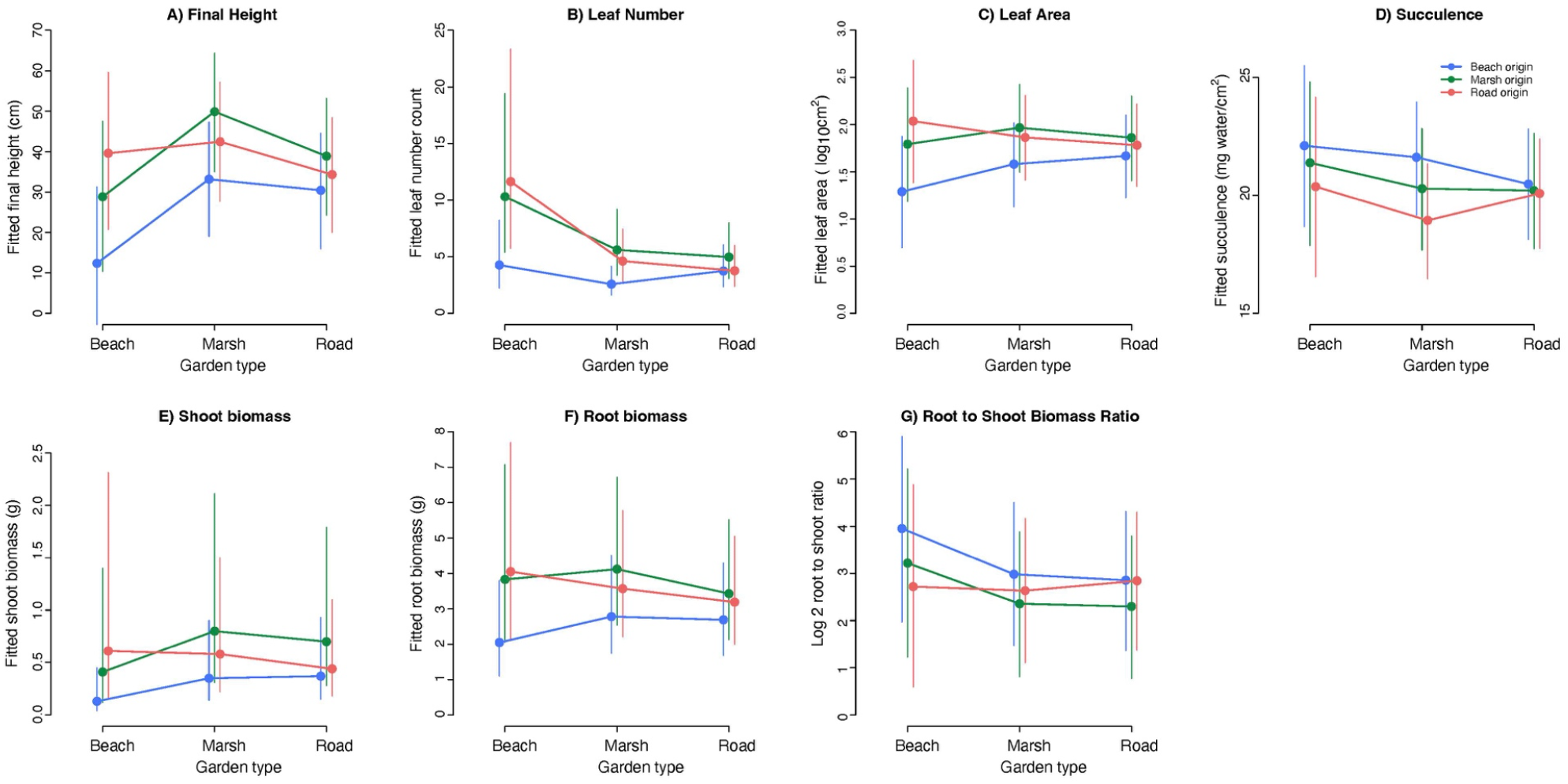
Reaction norms of (means +/− 95% CrI) across three transplant habitat gardens for plants from the three habitat origins: A) final height, B) total number of leaves, C) total leaf area of all leaves at final harvest, D) succulence as measured on all leaves at final harvest, E) dry shoot biomass, F) dry root biomass and G) dry root biomass to dry shoot biomass ratio at final harvest. Beach sites are depicted with blue lines, marsh sites with green and roadside sites with red.

**Table 2.** Tests for components of variance for each trait with random effects of origin site and transplant site, and fixed effects of source habitat type and transplant garden habitat type. The three tests of variance provide information about R^2^ of the full model versus just fixed effects (r2_nakagawa), R^2^ of the two random effects and combined fixed effects (rptR) and the contribution of each fixed effect without accounting for random effects (commonalityCoefficients). See methods for more details.

|  | **r2_nakagawa** |  | **rptR** |  |  | **commonalityCoefficients** |  |  |  |
| --- | --- | --- | --- | --- | --- | --- | --- | --- | --- |
|  | conditional<br>r2 (random<br>and fixed) | marginal r2<br>(fixed) | Origin site<br>(random) | Transplant site<br>(random) | Fixed effects | Unique to<br>Source type<br>(fixed) | Unique to<br>Garden type<br>(fixed) | Common<br>to Source<br>& Garden | Total |
| Final height | 0.448 | 0.12 | 0.021<br>[0, 0.073] | 0.421<br>[0.116, 0.612] | 0.077<br>[0.035, 0.35] | 0.127 | 0.063 | 0 | 0.186 |
| Total leaf number | 0.375 | 0.16 | NA | 0.143<br>[0, 0.212] | 0.08<br>[0.039, 0.236] | 0.048 | 0.072 | 0.008 | 0.128 |
| Total leaf area | 0.29 | 0.04 | 0.006<br>[0, 0.068] | 0.174<br>[0.011, 0.335] | 0.069<br>[0.03, 0.234] | 0.066 | 0.075 | 0.006 | 0.147 |
| Succulence | 0.185 | 0.03 | 0.019<br>[0, 0.095] | 0.204<br>[0, 0.379] | 0.018<br>[0.01, 0.166] | 0.021 | 0.008 | 0.0003 | 0.166 |
| Shoot biomass | 0.341 | 0.06 | NA | 0.367<br>[0.093, 0.561] | 0.045<br>[0.025, 0.258] | 0.093 | 0.023 | 0.002 | 0.119 |
| Root biomass | 0.366 | 0.05 | 0.016<br>[0, 0.077] | 0.423<br>[0.117, 0.620] | 0.027<br>[0.018, 0.299] | 0.088 | 0.062 | 0.007 | 0.157 |
| Total biomass | 0.379 | 0.06 | 0.013<br>[0, 0.07] | 0.427<br>[0.098, 0.606] | 0.034<br>[0.022, 0.283] | 0.099 | 0.051 | 0.007 | 0.157 |

In the beach gardens, plants originally from this habitat had only one-third the height, less than half the number of leaves, 30% less leaf area, one-fourth the shoot biomass and half as much root biomass as plants from the roadside habitats (Figure 1; Table S3). Plants from the beach habitat also had half as many leaves and nearly half the root biomass of plants from the marsh habitat when grown in the beach gardens. In the marsh gardens, plants from marsh habitats had two and half times the height and twice the number of leaves as plants from beach habitats. In addition, plants from beach habitats had greater succulence than plants from the roadside habitats (but not greater than plants from marsh habitats) when grown in the marsh gardens. In roadside garden, we found no differences among the groups for any of these traits. We also discovered that the responses of plants from marsh and roadside habitats were largely indistinguishable in any garden (Figures 1 and 2).

**Figure 2.**
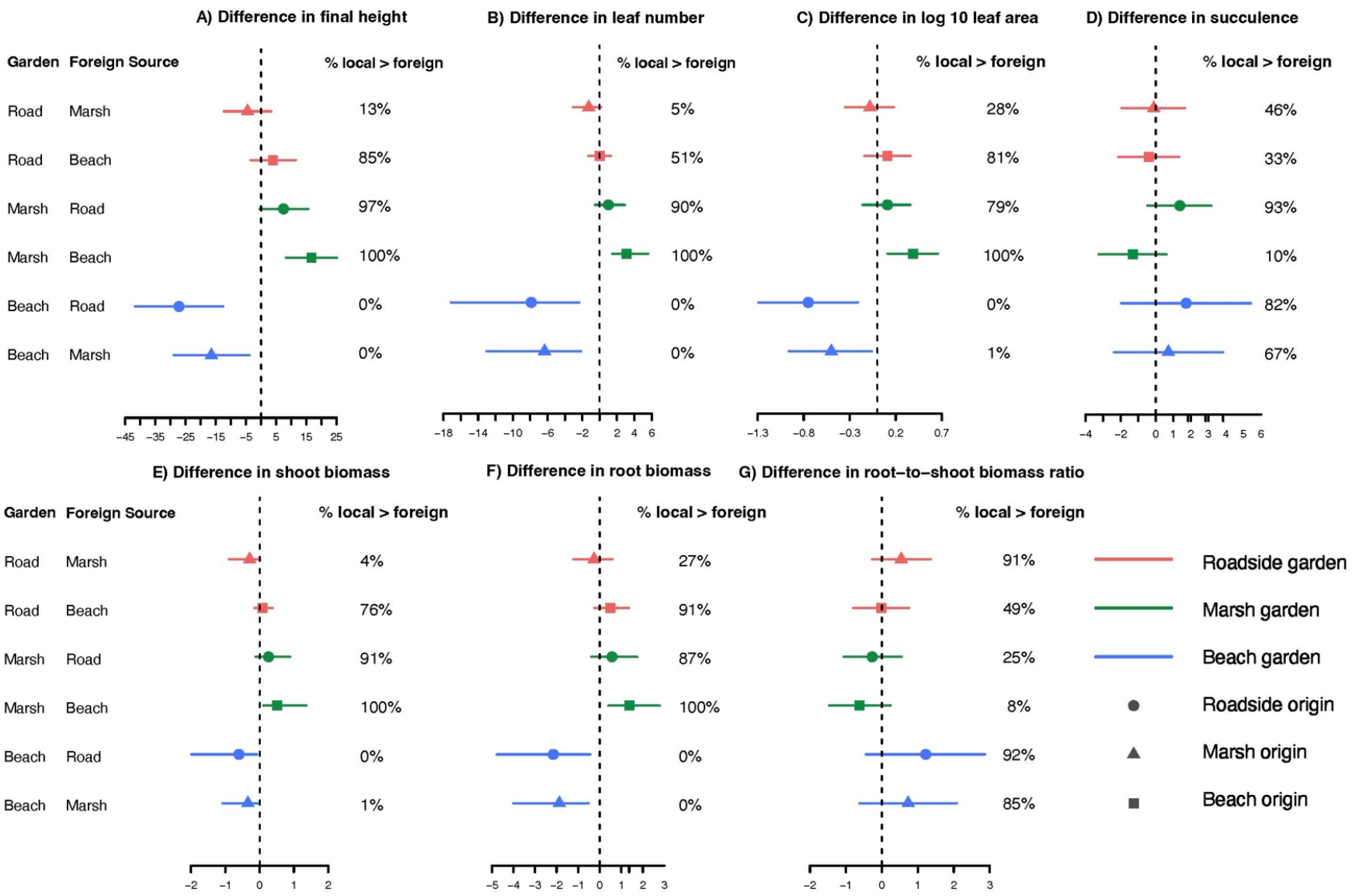
Differences in trait responses across three transplant habitat gardens for plants from the three habitat origins: A) final height, B) total number of leaves, C) total leaf area of all leaves at final harvest, D) succulence as measured on all leaves at final harvest, E) dry shoot biomass, F) dry root biomass and G) dry root biomass to dry shoot biomass ratio at final harvest. Beach sites are depicted with blue lines, marsh sites with green and roadside sites with red.

These findings were supported by examining simulated values of the differences between the groups of plants for each trait (Figure 2). In the beach habitat, beach plants were almost always shorter, had fewer leaves, had less leaf area, less root and shoot biomass than plants from marsh or roadside habitats (beach < marsh plants in 99 or 100% of the simulations). In the marsh garden, marsh plants were taller, had more leaves, greater leaf area, shoot and root biomass than beach plants in 100% of the simulations. Compared to roadside plants transplanted in the marsh garden, marsh plants were also usually taller (97% of the simulations), had more leaves (90%), greater leaf area (79%), shoot (91%) and root (87%) biomass but the differences in response were not as strong as comparisons to beach plants and did not meet our threshold for significance (Figure 1). In the roadside garden, again simulations did not support significant differences in pairwise comparisons of plants from different habitats.

### Explanatory power of the models of phenotypic variance –

Our linear mixed models explained 18 to 45% of the variation in the traits we measured (Table 2). However, we found that the majority of the variance was explained by the random effects (r2_nakagawa in Table 2) and in particular the “Transplant.site” which alone explained 14-43% of the variance in these traits. Only 2 to 8% of the variance was explained by the fixed effects of “Source type” or “Garden type” (rptR in Table 2) which were our main interest to test the general effects of habitats. Together the fixed effects were best able to explain variance in height and number of leaves (8%) and least predictive of succulence (2%). We used the commonalityCoefficients program to further examine the amount explained by each of the fixed effects. The source type explained twice as much of the variance as transplant garden type for height, succulence and total biomass, and three times the variance in shoot biomass, but less of the variance than transplant garden for the number of leaves. Source type explained a similar amount of the variance as transplant garden for leaf area.

When we evaluated the “Transplant group” as a fixed effect, the overall R^2^ changed very little (Table S2). On average the models changed by only 0.2%. The largest change in R^2^ was in the model for succulence which decreased from 29% in the original model without the effect of transplant group (Table 2) to 26% with the effect (Table S2). On average the Transplant group effect increased the amount of variance explained by fixed effects by 17%. Using commonalityCoefficients, we found that the unique contribution of transplant group was 29-65% of the variance explained by the combined fixed effects. For several traits (e.g., height, succulence, shoot root and total biomass) most of the variance explained by fixed effects was due uniquely to the transplant group effect (Table S2). Adding this effect also changed the amount of variance explained by “Source type” or “Garden type”. When the transplant group was included, the unique contribution of source type still explained twice as much of the variance as that of transplant garden type for shoot biomass, but a similar amount of variance as garden type for height, leaf area, succulence and total biomass. Using this model, source type explained half as much of the variance as transplant garden for the number of leaves.

### The effect of transplant habitats on fitness proxies –

We investigated the fitness proxies of total biomass (g) and survival. In the beach gardens, plants from beach habitats accumulated less biomass than plants from either marsh or roadside habitats in 100% of the simulations, contrary to predictions of local adaptation. On the other hand, in the marsh gardens, plants from the marsh habitat accumulated more biomass than plants from beaches (100% of the simulations) and tended to grow bigger than plants from roadsides (in 90% of the simulations but the effect size was smaller; Figure 3a). We found little support for differences in biomass among groups when grown in the roadside gardens.

**Figure 3.**
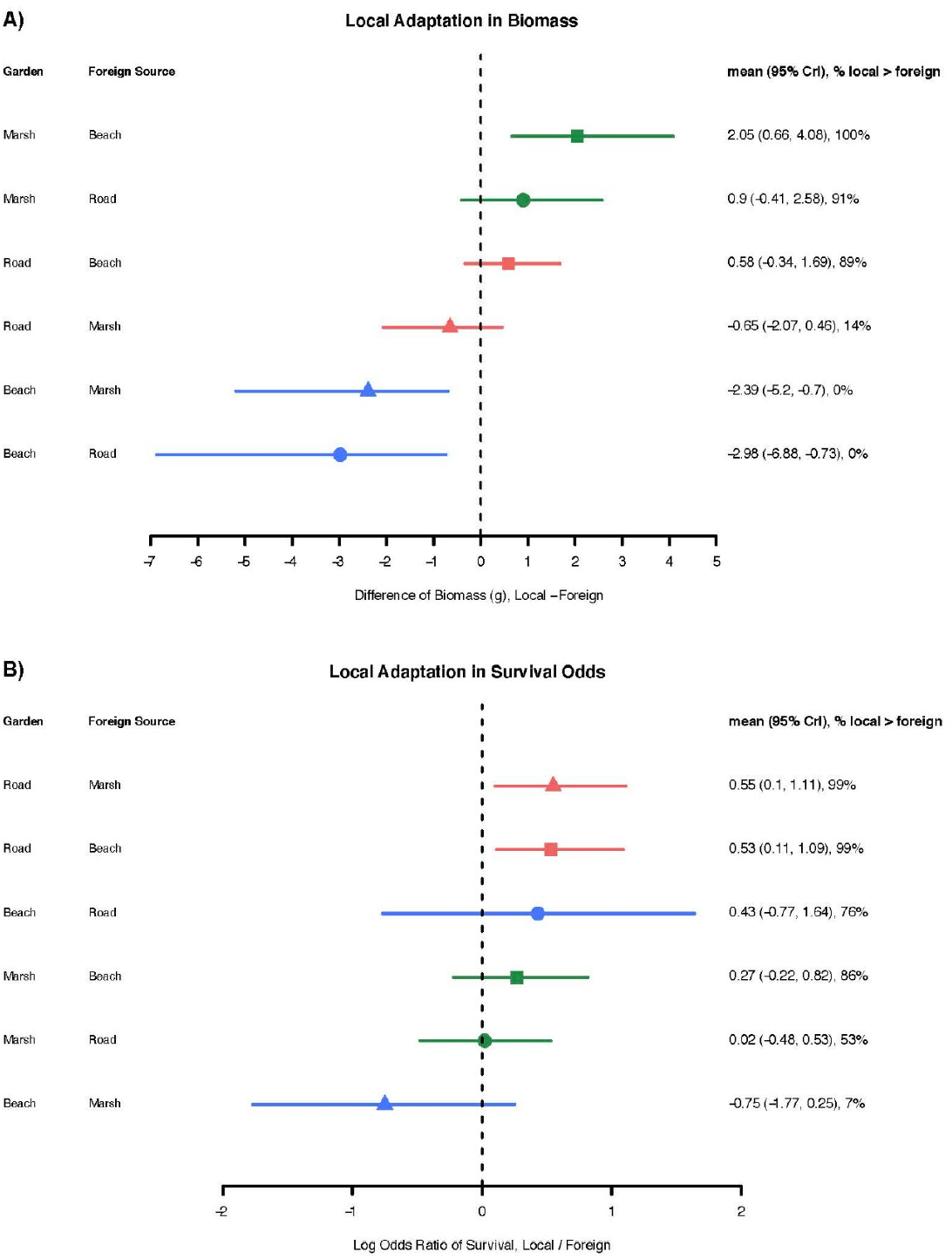
Local adaptation is supported in A) marsh plants compared to beach and roadside plants grown in marsh habitats as measured by total dry biomass (g; top two green lines) and B) survival of roadside plants compared to beach or marsh plants grown in roadside habitats (top two red lines). Symbols and whiskers are differences of fitted estimates and credible intervals estimated from statistical models (see Methods for details).

Our model explained approximately 38% of the variance in total biomass. This variance was largely determined by the random term “transplant garden site”: 43% of the variance was attributed to “transplant site” when “transplant group” was not included (Table 2), 36% when “transplant group” was included as a fixed effect (Table S2). The fixed effect of origin habitat type explained almost twice as much as that of transplant garden habitat type, but combined they explained only 6% of the variance in biomass (Table 2). When transplant group is included as a fixed effect, the R^2^ jumps to 24% explained by combined fixed effects (according to results of r2_nakagawa, Table S2) and the effect of origin habitat type still explains more than that of transplant garden habitat type (19% compared to 13% of the variance due to fixed effects which translates to approximately 4% and 3% of the overall variance in this model).

Mortality was high across the experiment, but particularly in the beach habitat garden sites (average 89% mortality; Table 3), where one site was completely washed away in a storm (PJB) and two other sites suffered 95-97% mortality. Plants from marsh and roadside habitats showed an average of 5.2 – to 14.2 – fold decrease of survival odds when transplanted into beach habitats (probability 0.98-1; Figure 4.). Plants from the marsh habitats suffered less in survival odds than roadside plants (Log Odds Ratio 2.24 vs 3.8). In contrast, the beach plants had an average 7-fold increase in survival odds when transplanted to either of the other habitats (Figure 4). The survival odds in reciprocal transplants between road and marsh habitats were similar. In sum, the survival data further suggested that the beach habitat was the most challenging environment out of the three tested, leading to reduced odds of survival for plants from all three habitats. Meanwhile, marsh and roadside habitats were similar.

**Table 3.**
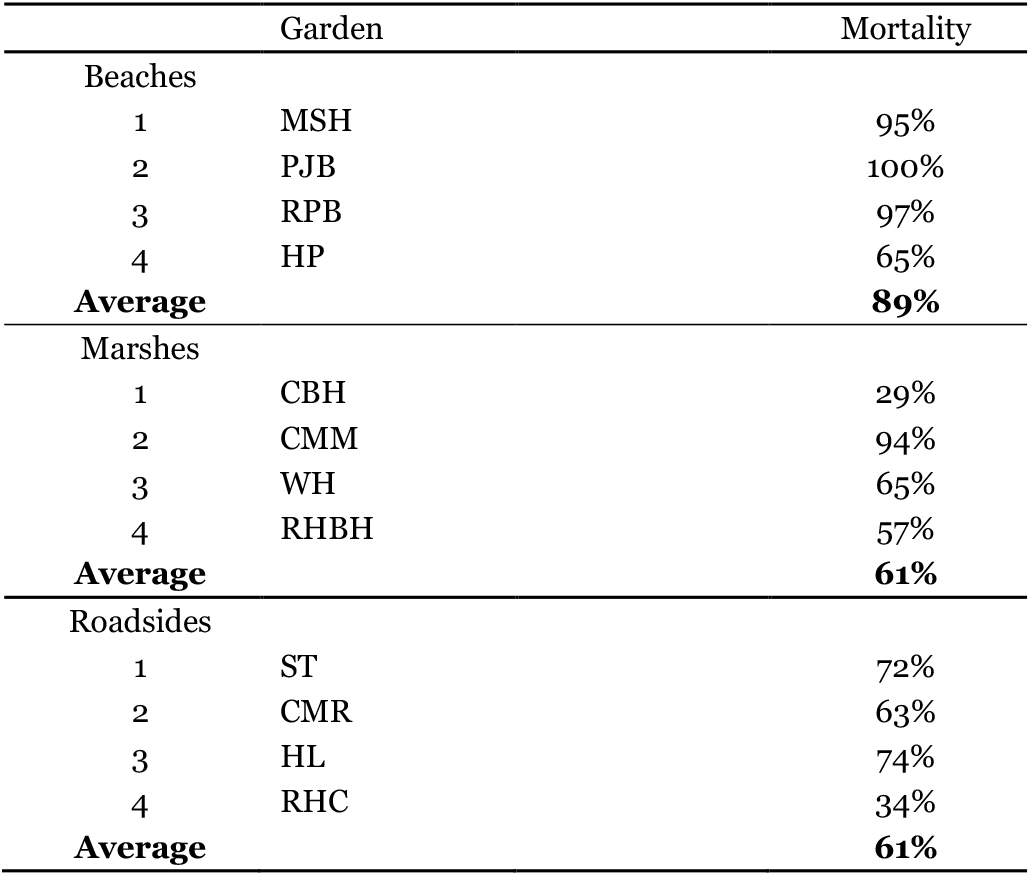
Breakdown of mortality by group within transplant habitat and average for each habitat type

**Figure 4.**
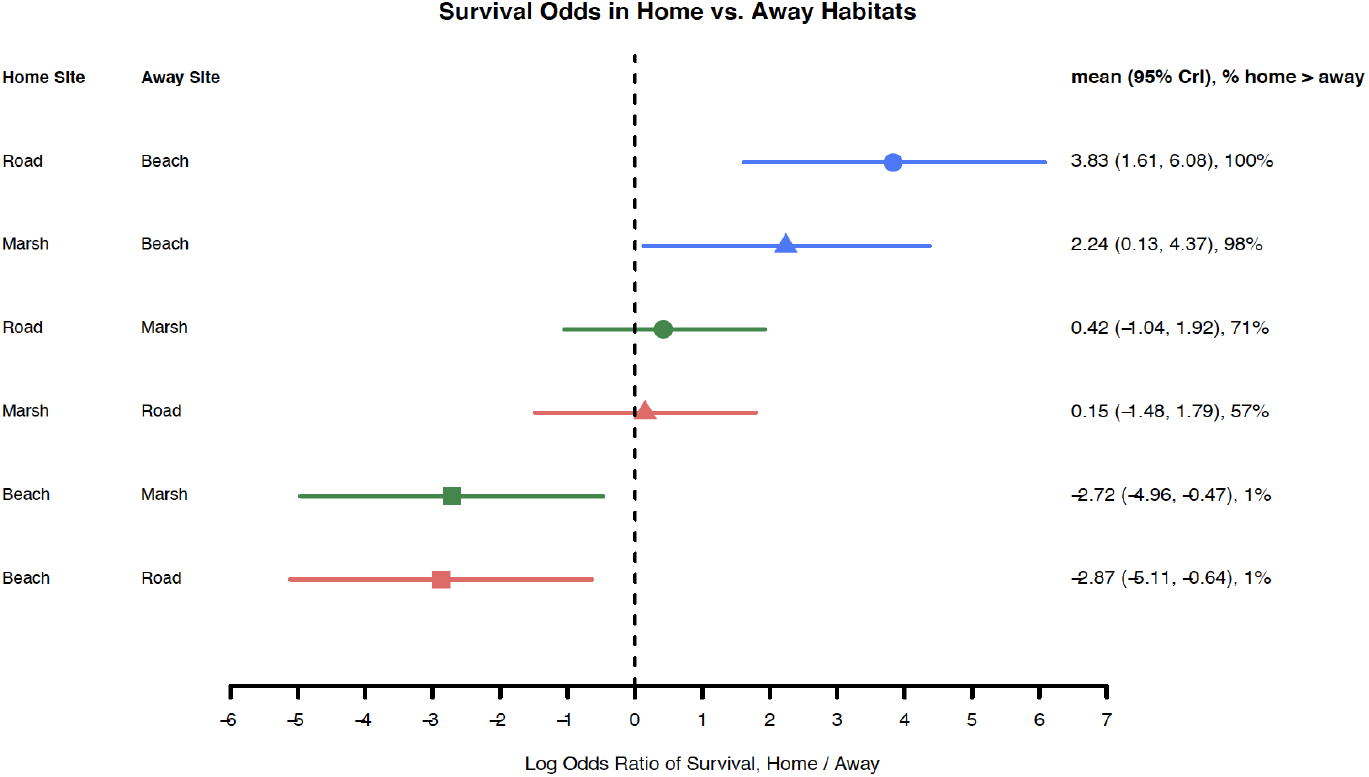
Evidence of differences in survival across habitats. Marsh and roadside plants grow better in their home compared to beach sites while beach plants grow better in marsh and roadside habitats than their home habitat (see Methods for details). Beach sites are depicted with blue lines, marsh sites with green and roadside sites with red.

We found no support for local adaptation in plants from beach habitats or marsh habitats when comparing survival in their home habitat to foreign plants in that garden (Figure 3b). However, plants from roadside habitats showed better survival when compared to foreign plants from the beach or the marsh (Figure 3b).

## DISCUSSION

In this study, we investigated how one of the world’s most invasive plants may be adapting to three different habitats on Long Island, NY. Many studies have demonstrated that significant differences in habitat characteristics can result in adaptive differentiation within species, even under high levels of gene flow between habitats (Antonovics and Bradshaw, 1970; Linhart and Grant, 1996; Sambatti and Rice, 2006; Papadopulos et al., 2021; Zerebecki et al., 2021 but see Leimu and Fischer, 2008). Introduced species in particular have been highlighted because they can evolve rapidly in response to novel conditions (Lee, 2002; Leger and Rice, 2007; Dlugosch and Parker, 2008). We took advantage of replicate populations to test the generality of adaptation to different habitats. Despite the wide variation among sites within habitat types, we revealed differentiation between plants from beach, marsh and roadside populations for most of these purportedly adaptive traits, as well as for fitness. In addition, we found some support for local adaptation.

### Phenotypic plasticity in Japanese knotweed –

Overall, the variation in phenotypes in our study was most often best explained by the local conditions of the transplant garden site. These findings indicate the importance of plasticity, which has often been highlighted in invasion ecology (Bossdorf et al., 2005, 2008; Freeman and Byers, 2006; Richards et al., 2006; Geng et al., 2007; Muth and Pigliucci, 2007; Funk, 2008; Loomis and Fishman, 2009; Walls, 2010). In our previous greenhouse study, plants from both roadside and marsh habitats were also highly plastic in response to treatment with salt, and within sites there were significant differences in most traits and trait plasticity.

### Habitat differences –

In order for adaptive differentiation to occur, habitats must select for different trait means, plasticities or relationships among traits. The beach habitat was typically open with plenty of sun. In contrast, the marsh habitats were typically under a canopy of tall (2.5-3 m), well established knotweed or *Phragmites* plants, and the roadside sites were typically under a canopy of trees. Our pairwise comparisons of plants from different habitats across transplant gardens indicated that plants were significantly different for every trait except R:S. Most of the differences were manifest in the beach common gardens where beach plants were significantly shorter, with fewer leaves, and less biomass than plants from marsh and roadside sites. This finding of more differences in the beach habitat is similar to our previous study where we found differences in succulence and root to shoot biomass ratio only under salt addition, but not under pure water conditions (Richards et al., 2008; see similar results in *Borrichia frutescens* Richards et al., 2010). Similarly, a recent reciprocal transplant study in the salt marsh cordgrass *Spartina alterniflora* identified more extreme differences in the “Tall Spartina zone” habitat for survival, maximum height, root to shoot biomass ratio and total biomass (Zerebecki et al., 2021). In these examples, the conditions that were more challenging for the plants also elicited more differences between plants.

We did find a few differences elicited in the marsh transplant gardens where marsh plants tended to be larger than the others. We expected that succulence could be important for invasion of the saline marsh habitat because the ability to become succulent, and dilute the toxic effect of concentrated salt ions, is essential for many species in salt environments (Flowers et al., 1977). For example, *Salsola kali* originating from different habitats was found to have dramatic intraspecific variation in succulence and the salt tolerant subspecies *S. kali traga* was able to increase succulence more than the non-salt tolerant *S. kali ruthenica* (Reimann and Breckle, 1995). However, we did not find consistent response in succulence in our previous work with these knotweed taxa. Instead, we found a lot of variation among knotweed genets for succulence in response to salt treatments, including several genets that seemed to display no change in succulence (Richards et al., 2008). In this study, the only difference in succulence we found was that the beach plants were more succulent than the roadside plants in the marsh transplant garden. Succulence could aid in the adaptation to saline habitats in *Reynoutria,* but could be specific to certain genets or conditions that we did not explore with our current design. The plants with by far the highest amount of succulence in this study were those from the Marsh 3 site grown in their home marsh environment and plants from the Marsh 1 site and Beach 1 site grown at the beach. This increased level of succulence may have been an important trait contributing to the slight advantage in biomass exhibited by Marsh 3 plants at the marsh and the increased survival of Marsh 1 and Beach 1 plants at the beach where plants from the roadside did not survive at all.

The differences in phenotype elicited by these habitats could result in adaptive differentiation if there is heritable variation for these traits within the populations. We did not detect any genet-level variation within populations in this study. However, our power to detect this level of variation was limited by the high mortality. The random effect of source site did not explain much of the variation, but the source habitat type was a better predictor of variation than transplant garden habitat type for most traits.

### Signature of local adaptation –

We compared performance of local plants with that of the foreign plants for fitness proxies (total biomass and survival) in each of the common gardens to assess local adaptation. Previously, a similar reciprocal transplant study in knotweed found little support for local adaptation along a latitudinal gradient in three populations from similar temperate deciduous forest habitats (VanWallendael et al., 2018). Comparing responses to different habitat types on a more local scale, we found support for local adaptation in plants from the marsh: they out-performed plants from both beach and roadside habitats in biomass. This is surprising given the previous work in the greenhouse which did not support adaptive response to salt treatments. In the current study, the salt marsh plants were the only plants that accumulated more biomass in their home site. This growth advantage, however, did not translate into advantage in survival of marsh plants over foreign plants, in the marsh garden. Instead, there were no differences in survival due to habitat of origin in the marsh transplant garden.

Plants from beach and roadside habitats failed to accumulate more total biomass in their home environment compared to foreign plants. Beach plants tended to allocate more resources to roots, but such allocation did not increase their odds of survival compared to foreign plants. In fact, beach plants seemed to be mal-adapted, at least during the time frame of our study. It is unclear whether preferential allocation to roots could eventually lead to an advantage, or if this response is constrained by other factors, or could have been detected in a longer-term study. Plants from the beach habitat have a greater probability to grow larger in either of the away habitats. Growing on the beach on average reduced the biomass accumulation by 1.03-1.1 grams, indicating that beach habitats are suboptimal for plant growth.

We also found support for local adaptation among roadside plants, which were better able to survive in their home sites. Roadside plants had the best survival odds when compared to foreign plants. This is somewhat surprising considering that plants from roadside habitats are largely indistinguishable from plants from marsh habitats for most traits that we measured.

### Sources of phenotypic differentiation –

In our previous work, we used cytology and AFLP markers to show that most of these populations consist of a few *R. × bohemica* hybrids (Richards et al., 2008, 2012). Plants from both roadside and marsh habitats were highly plastic in response to treatment with salt in the greenhouse, and even though clonal diversity was low in these populations, within sites there were significant differences in most traits and trait plasticities. Several studies have demonstrated that hybridization can result in significant changes in trait expression (e.g., transgressive traits) with important ecological consequences (Gaskin and Schaal, 2002; Rosenthal et al., 2002, 2008; Lexer et al., 2003; Johnston et al., 2004; Karrenberg et al., 2006; Parepa et al., 2014). In fact, novel traits resulting from hybridization are considered an important feature that allows expansion into novel habitats in several systems (Ellstrand and Schierenbeck, 2000; Lexer et al., 2003; Johnston et al., 2004; Karrenberg et al., 2006). For example, hybrids between the invasive *Carpobrotus edulis* and the native *C. chilensis* have higher biomass in response to low salinity treatments under low nutrient conditions (Weber and D’Antonio, 1999), indicating that they may have an advantage in nutrient poor soils. These and other studies suggest that recombination of different traits may allow for rapid adaptation to new environments (Anderson and Stebbins, 1954; Ellstrand and Schierenbeck, 2000; Facon et al., 2005; Gross and Rieseberg, 2005).

Hybridization between *R. japonica* and *R. sachalinensis* to form *R. × bohemica* has also been considered an important mechanism in the Japanese knotweed *s.l* invasion (Pysek et al., 2003; Bímová et al., 2004; Mandák et al., 2005; Bailey et al., 2009). We considered that transgressive trait segregation. might therefore be an important contributor to success in this diversity of habitats and that we would find particularly aggressive *R. × bohemica* genotypes. We found that a few of our populations were characterized as *R. japonica* (Marsh 2, Roadside 2 and Beach 4). According to ANOVA they did not respond differently to these habitats than *R. × bohemica*. However, the ANOVA was performed only on surviving individuals and a logistic regression of taxon on survival suggests that taxon alone explains a small, but significant amount of the variation in survival. For the long-term success of *R. japonica,* it is meaningful that plants from these populations did not *survive* as well as the *R. × bohemica* hybrids in our field transplants. Plants from Marsh 2 and Roadside 2, for instance, did not survive at the beach and marsh transplants had low survival on the roadside. This suggests that the hybrids have an increased ability to establish in the most diverse and stressful habitats. In combination, these studies show that complex ecologically relevant environments elicit differences in phenotype that are not detectable by manipulation of salinity alone under controlled conditions, which further underscores the importance of conducting studies in the field (Endler, 1986; Kingsolver et al., 2001; Kawecki and Ebert, 2004; Sambatti and Rice, 2006; Leimu and Fischer, 2008).

Even considering that the few genotypes that have invaded these habitats may have benefited from transgressive traits, the dramatically varied response to the transplant habitats from what should only be a few genotypes is surprising. For example, the plants from Marsh 4 and Roadside 4 were identical across AFLP markers (Richards et al., 2012), and they have almost identical survival at the roadside site. However, at the marsh transplant the marsh plants had a significantly higher survival rate. Under different circumstances, this could reflect the importance of maternal effects or provisioning. However, in our study we took care to start plants with similar initial rhizome weights.

A potentially important possibility is that persistent epigenetic effects may have resulted from the hybridization process or may have been induced by exposure to these dramatically different environments. We have reported a surprising level of epigenetic variation in these populations compared to levels of sequence-based variation found with AFLP (Richards et al., 2012). Epigenetic effects have been suggested as a source of phenotypic variation in ecologically relevant traits, but they have not yet been explored extensively in studies of invasive species (Mounger et al., 2021; Hawes et al. 2018). In some cases, environmentally-induced epigenetic changes may be inherited by future generations (Jablonka and Raz, 2009; Verhoeven et al., 2010; Richards et al., 2017; Bonduriansky and Day, 2018; Richards and Pigliucci, 2020) and therefore could contribute to explaining short-term adaptation to novel environments. Moreover, epigenetic processes are an important component of hybridization events (Rapp and Wendel, 2005; Salmon et al., 2005; Flowers and Burton, 2006).

## CONCLUSIONS

Although we found only limited support for local adaptation, this is not so unusual (VanWallendael et al., 2018). Leimu & Fischer (2008) reported a meta-analysis of local adaptation studies where they found that plants from “home” populations outperformed the “foreign” plants in both habitat types in only 51% of the studies surveyed. These findings were independent of plant longevity, mating system, clonality or habitat type and the authors concluded that local adaptation may not be as common as it is assumed. Considering the potentially random sampling of genotypes during the invasion process, genetic drift may play a large role in shaping the evolutionary trajectory of these populations (Keller and Taylor, 2008; Prentis et al., 2008). Identifying adaptive changes in a small founding population is difficult because it requires identifying the source of the invasion and comparing responses of the invaders to those of the source material (Bossdorf et al., 2008; Keller and Taylor, 2008; Prentis et al., 2008; Colautti and Lau, 2015); e.g. (Bock et al., 2018; Exposito-Alonso et al., 2018).

Despite our mixed evidence, adaptive processes could still be important for most of the populations, since transplants maintained biomass across at least two if not all three habitats. The current study confirms our findings from the greenhouse that there is phenotypic differentiation among these populations of Japanese knotweed, some of which is attributed to their source habitat. Some of the plasticity in these traits and in fitness are likely to be passive responses to resource limitation and stress, but “active” or adaptive plasticity in underlying morphological and physiological traits may help to minimize the fitness loss in these environments (van Kleunen and Fischer, 2005; Van Kleunen and Fischer, 2007; Murren et al., 2015; Bock et al., 2018). Whatever the mechanisms of divergence, which could include drift, selection, transgressive segregation, nongenetic effects, and genetic accommodation, this study demonstrates that there is persistent phenotypic variation present in the populations of interest. This variation in ecologically important traits provides the potential for future adaptation that could increase the already high rate of spread of this species complex in North America, and in salt marsh and beach habitats in particular. Understanding the components that contribute to the success of this extensively clonal plant with little or no genetic variation could require reevaluating how we measure adaptation.

## Acknowledgments

The authors thank Ramona Walls for valuable assistance in finding field sites and making connections with local government agencies and organizations. We also thank Larry Gottschamer, Kristi Adams, Oliver Bossdorf, Tara George, Norris Muth and Radha Parameswaran for assistance in the field and with harvesting as well as Stony Brook University greenhouse staff Mike Axelrod and John Clumpp. We thank Fränzi Korner-Nievergelt, Ramona Irimia, Madalin Parepa, Robert Rauschkolb and Oliver Bossdorf for critical advice on data analysis. This work was supported by the Research Foundation of the State University of New York (to MP), New York SEA Grant (to MP and CLR), and the Federal Ministry of Education and Research (BMBF; MOPGA Project ID 306055 to CLR).

## Author Contributions

C.L.R. and M.P. designed the study. C.L.R. implemented and maintained the experiment and collected all data. W.Y. and C.L.R. completed statistical analyses. C.L.R. wrote the manuscript and all authors contributed to the writing.

## Data Availability Statement

Data and code for data analysis have been submitted for review on the Dryad Digital Repository: https://doi.org/10.5061/dryad.wdbrv15qz

---

**Table S1.**
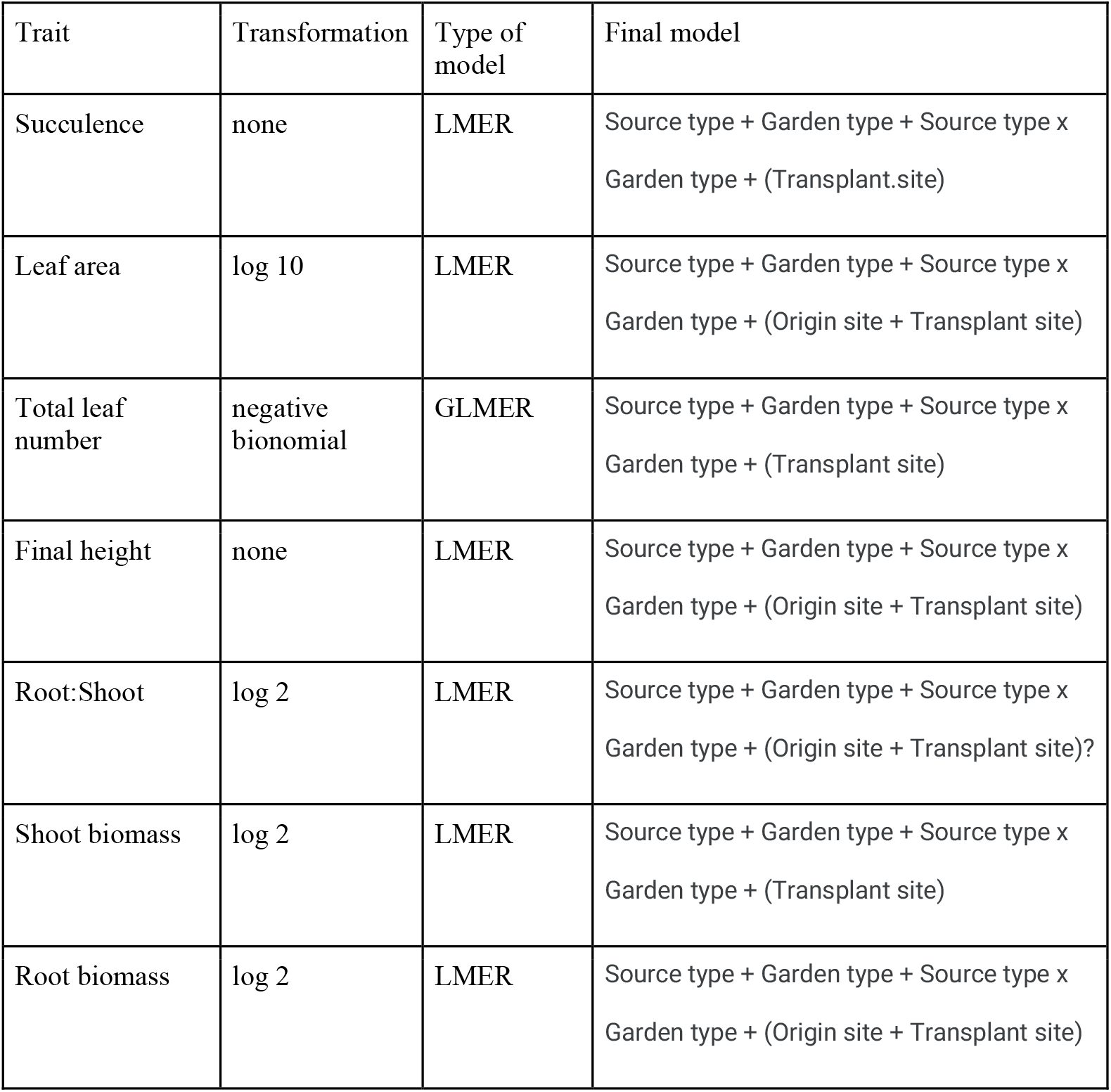
Final models for the seven traits.

| Trait | Transformation | Type of model | Final model |
| --- | --- | --- | --- |
| Succulence | none | LMER | Source type + Garden type + Source type x Garden type + (Transplant.site) |
| Leaf area | log 10 | LMER | Source type + Garden type + Source type x Garden type + (Origin site + Transplant site) |
| Total leaf number | negative bionomial | GLMER | Source type + Garden type + Source type x Garden type + (Transplant site) |
| Final height | none | LMER | Source type + Garden type + Source type x Garden type + (Origin site + Transplant site) |
| Root:Shoot | log 2 | LMER | Source type + Garden type + Source type x Garden type + (Origin site + Transplant site)? |
| Shoot biomass | log 2 | LMER | Source type + Garden type + Source type x Garden type + (Transplant site) |
| Root biomass | log 2 | LMER | Source type + Garden type + Source type x Garden type + (Origin site + Transplant site) |

**Table S2.** As with table 2 in the main text, we provide tests for components of variance for each trait with random effects of origin site and transplant site, and fixed effects of source habitat type and transplant garden habitat type as well as the fixed effect of transplant group. The three test of variance provide information about R^2^ of the full model versus just fixed effects (r2_nakagawa), R^2^ of the two random effects and combined fixed effects (rptR) and the contribution of each fixed effect without accounting for random effects (commonalityCoefficients). See methods for more details.

| | $r2\_nakagawa$ | | $rptR$ | | |
| --- | --- | --- | --- | --- | --- |
|  | conditional r2 (random and fixed) | marginal r2 (fixed) | Origin site (random) [CI] | Transplant site (random) [CI] | Fixed effects [CI] |
| Final height (396) | 0.466 | 0.334 | 0.019<br>[0, 0.072] | 0.3<br>[0.032, 0.497] | 0.24<br>[0.135, 0.563] |
| Total leaf number (395) | 0.374 | 0.364 | NA | 0<br>[0, 0] | 0.235<br>[0.184, 0.321] |
| Total leaf area (395) | 0.264 | 0.231 | 0.009<br>[0, 0.06] | 0.165<br>[0, 0.362] | 0.136<br>[0.077, 0.365] |
| Succulence (372) | 0.185 | 0.067 | 0.015<br>[0, 0.095] | 0.253<br>[0.002, 0.074] | 0.045<br>[0.038, 0.334] |
| Shoot biomass (395) | 0.334 | 0.275 | NA | 0.208<br>[0.005, 0.414] | 0.226<br>[0.116, 0.483] |
| Root biomass (395) | 0.382 | 0.209 | 0.015<br>[0, 0.064] | 0.379<br>[0.045, 0.588] | 0.148<br>[0.071, 0.528] |
| Total biomass (395) | 0.394 | 0.239 | 0.013<br>[0, 0.057] | 0.361<br>[0.027, 0.056] | 0.169<br>[0.086, 0.535] |

|  | **commonalityCoefficients** |  |  |  |  |  |  | Total |
| --- | --- | --- | --- | --- | --- | --- | --- | --- |
|  | Unique to Transplant group (fixed) | Unique to Source type (fixed) | Unique to Garden type (fixed) | Common to Group & Source type | Common to Group & Garden type | Common to Source type & Garden type | Common to Group, Source & Garden type |  |
| Final height | 0.175 | 0.055 | 0.062 | 0.0713 | 0.0003 | 0.003 | -0.006 | 0.359 |
| Total leaf number | 0.075 | 0.039 | 0.076 | 0.0091 | -0.0046 | -0.002 | 0.0097 | 0.202 |
| Total leaf area | 0.060 | 0.042 | 0.056 | 0.0235 | 0.0189 | -0.002 | 0.0073 | 0.147 |
| Succulence | 0.053 | 0.012 | 0.011 | 0.009 | -0.0038 | 0.002 | -0.002 | 0.029 |
| Shoot biomass | 0.155 | 0.042 | 0.017 | 0.0507 | 0.0075 | 0.001 | 0.0013 | 0.119 |
| Root biomass | 0.103 | 0.048 | 0.047 | 0.0402 | 0.0156 | -0.001 | 0.0081 | 0.157 |
| Total biomass | 0.121 | 0.052 | 0.036 | 0.0477 | 0.0148 | -0.0004 | 0.0072 | 0.157 |

## Literature Cited

Allendorf, F. W., and L. L. Lundquist. 2003. Introduction: Population biology, evolution, and control of invasive species. Conservation biology: the journal of the Society for Conservation Biology 17: 24–30.

Anderson, E., and G. L. Stebbins. 1954. Hybridization as an Evolutionary Stimulus. Evolution; international journal of organic evolution 8: 378–388.

Antonovics, J., and A. D. Bradshaw. 1970. Evolution in closely adjacent plant populations VIII. Clinal patterns at a mine boundary. Heredity 25: 349–362.

Bailey, J. P. 2013. The Japanese knotweed invasion viewed as a vast unintentional hybridisation experiment. Heredity 110: 105–110.

Bailey, J. P., K. Bímová, and B. Mandák. 2009. Asexual spread versus sexual reproduction and evolution in Japanese Knotweed s.l. sets the stage for the ‘Battle of the Clones’. Biological invasions 11: 1189–1203.

Bailey, J. P., and A. P. Conolly. 2000. Prize-winners to pariahs - a history of Japanese knotweed s.l. (Polygonaceae) in the British Isles. Watsonia 23: 93–110.

Bailey, J. P., and R. Wisskirchen. 2004. The distribution and origins of Faúopia × bohemica (Polygonaceae) in Europe. Nordic Journal of Botany 24: 173–199.

Baker, H. G. 1965. Characteristics and modes of origin of weeds. In H. G. Baker, and G. L. Stebbins [eds.], The genetics of colonizing species, 147–172. Academic Press Inc., NY.

Barney, J. N. 2006. North American History of Two Invasive Plant Species: Phytogeographic Distribution, Dispersal Vectors, and Multiple Introductions. Biological invasions 8: 703–717.

Bates, D., M. Mächler, B. Bolker, and S. Walker. 2015. Fitting linear mixed-effects models Usinglme4. Journal of statistical software 67.

Bímová, K., B. Mandák, and I. Kašparová. 2004. How does Reynoutria invasion fit the various theories of invasibility? Journal of vegetation science: official organ of the International Association for Vegetation Science 15: 495–504.

Bímová, K., B. Mandák, and P. Pyšek. 2001. Experimental control of Reynoutria congeners: a comparative study of a hybrid and its parents. Plant invasions: species ecology and ecosystem management, 283–290. Backhuys Publishers.

Bock, D. G., C. Caseys, R. D. Cousens, M. A. Hahn, S. M. Heredia, S. Hübner, K. G. Turner, et al. 2015. What we still don’t know about invasion genetics. Molecular ecology 24: 2277–2297.

Bock, D. G., M. B. Kantar, C. Caseys, R. Matthey-Doret, and L. H. Rieseberg. 2018. Evolution of invasiveness by genetic accommodation. Nature ecology & evolution 2: 991–999.

Bolker, B. M., M. E. Brooks, C. J. Clark, S. W. Geange, J. R. Poulsen, M. H. H. Stevens, and J.-S. S. White. 2009. Generalized linear mixed models: a practical guide for ecology and evolution. Trends in ecology & evolution 24: 127–135.

Bonduriansky, R., and T. Day. 2018. Extended heredity. Princeton University Press, Princeton, NJ.

Bossdorf, O., H. Auge, L. Lafuma, W. E. Rogers, E. Siemann, and D. Prati. 2005. Phenotypic and genetic differentiation between native and introduced plant populations. Oecologia 144: 1–11.

Bossdorf, O., A. Lipowsky, and D. Prati. 2008. Selection of preadapted populations allowed Senecio inaequidens to invade Central Europe. Diversity & distributions 14: 676–685.

Bucharova, A., W. Durka, N. Hölzel, J. Kollmann, S. Michalski, and O. Bossdorf. 2017. Are local plants the best for ecosystem restoration? It depends on how you analyze the data. Ecology and evolution 7: 10683–10689.

Bucharova, A., M. Frenzel, K. Mody, M. Parepa, W. Durka, and O. Bossdorf. 2016. Plant ecotype affects interacting organisms across multiple trophic levels. Basic and applied ecology 17: 688–695.

Cheplick, G. P. 2006. A modular approach to biomass allocation in an invasive annual (Microstegium vimineum; Poaceae). American journal of botany 93: 539–545.

Colautti, R. I., and J. A. Lau. 2015. Contemporary evolution during invasion: evidence for differentiation, natural selection, and local adaptation. Molecular ecology 24: 1999–2017.

Daehler, C., and D. Strong. 1997. Hybridization between introduced smooth cordgrass (Spartina alterniflora; Poaceae) and native California cordgrass (S. foliosa) in San Francisco Bay, California, USA. American journal of botany 84: 607.

Davidson, A. M., M. Jennions, and A. B. Nicotra. 2011. Do invasive species show higher phenotypic plasticity than native species and, if so, is it adaptive? A meta-analysis: Invasive species have higher phenotypic plasticity. Ecology letters 14: 419–431.

Del Tredici, P. 2017. The introduction of Japanese knotweed, Reynoutria japonica, into North America. The Journal of the Torrey Botanical Society 144: 406–416.

Dlugosch, K. M., and I. M. Parker. 2008. Invading populations of an ornamental shrub show rapid life history evolution despite genetic bottlenecks. Ecology letters 11: 701–709.

Dlugosch, K. M., and I. M. Parker. 2007. Molecular and quantitative trait variation across the native range of the invasive species Hypericum canariense: evidence for ancient patterns of colonization via pre-adaptation? Molecular ecology 16: 4269–4283.

Donovan, L. A., S. A. Dudley, D. M. Rosenthal, and F. Ludwig. 2007. Phenotypic selection on leaf water use efficiency and related ecophysiological traits for natural populations of desert sunflowers. Oecologia 152: 13–25.

Donovan, L. A., J. H. Richards, and M. W. Muller. 1996. Water relations and leaf chemistry ofChrysothamnus nauseosusssp.consimilis(Asteraceae) andSarcobatus vermiculatus(Chenopodiaceae). American journal of botany 83: 1637–1646.

Doroszuk, A., M. W. Wojewodzic, G. Gort, and J. E. Kammenga. 2008. Rapid divergence of genetic variance-covariance matrix within a natural population. The American naturalist 171: 291–304.

Dudley, S. A. 1996a. Differing selection on plant physiological traits in response to environmental water availability: a test of adaptive hypotheses. Evolution; international journal of organic evolution 50: 92–102.

Dudley, S. A. 1996b. The response to differing selection on plant physiological traits: evidence for local adaptation. Evolution; international journal of organic evolution 50: 103–110.

Durka, W., O. Bossdorf, D. Prati, and H. Auge. 2005. Molecular evidence for multiple introductions of garlic mustard (Alliaria petiolata, Brassicaceae) to North America. Molecular ecology 14: 1697–1706.

Ellstrand, N. C., and K. A. Schierenbeck. 2000. Hybridization as a stimulus for the evolution of invasiveness in plants? Proceedings of the National Academy of Sciences of the United States of America 97: 7043–7050.

Endler, J. A. 1986. Natural Selection in the Wild. Princeton University Press.

Exposito-Alonso, M., C. Becker, V. J. Schuenemann, E. Reiter, C. Setzer, R. Slovak, B. Brachi, et al. 2018. The rate and potential relevance of new mutations in a colonizing plant lineage. PLoS genetics 14:e1007155.

Facon, B., P. Jarne, J. P. Pointier, and P. David. 2005. Hybridization and invasiveness in the freshwater snail Melanoides tuberculata: hybrid vigour is more important than increase in genetic variance. Journal of evolutionary biology 18: 524–535.

Flowers, J. M., and R. S. Burton. 2006. Ribosomal RNA Gene Silencing in Interpopulation Hybrids of Tigriopus californicus: Nucleolar Dominance in the Absence of Intergenic Spacer Subrepeats. Genetics 173: 1479–1486.

Flowers, T. J., P. F. Troke, and A. R. Yeo. 1977. The mechanism of salt tolerance in halophytes. Annual review of plant physiology 28: 89–121.

Forman, J., and R. V. Kesseli. 2003. Sexual reproduction in the invasive species Fallopia japonica (Polygonaceae). American journal of botany 90: 586–592.

Franks, S. J., and A. E. Weis. 2008. A change in climate causes rapid evolution of multiple life-history traits and their interactions in an annual plant. Journal of evolutionary biology.

Freeman, A. S., and J. E. Byers. 2006. Divergent induced responses to an invasive predator in marine mussel populations. Science 313: 831–833.

Funk, J. L. 2008. Differences in plasticity between invasive and native plants from a low resource environment. The Journal of ecology 96: 1162–1173.

Gammon, M. A., J. L. Grimsby, D. Tsirelson, and R. Kesseli. 2007. Molecular and morphological evidence reveals introgression in swarms of the invasive taxa Fallopia japonica, F. sachalinensis, and F. xbohemica (Polygonaceae) in the United States. American journal of botany 94: 948–956.

Gammon, M. A., and R. Kesseli. 2010. Haplotypes of Fallopia introduced into the US. Biological invasions 12: 421–427.

Gaskin, J. F., and B. A. Schaal. 2002. Hybrid Tamarix widespread in U.S. invasion and undetected in native Asian range. Proceedings of the National Academy of Sciences of the United States of America 99: 11256–11259.

Gaskin, J. F., M. Schwarzländer, F. S. Grevstad, M. A. Haverhals, R. S. Bourchier, and T. W. Miller. 2014. Extreme differences in population structure and genetic diversity for three invasive congeners: knotweeds in western North America. Biological invasions 16: 2127–2136.

Gelman, A., and Y.-S. Su. 2020. arm: Data Analysis Using Regression and Multilevel/Hierarchical Models. R package version 1. 11–2.

Geng, Y.-P., X.-Y. Pan, C.-Y. Xu, W.-J. Zhang, B. Li, J.-K. Chen, B.-R. Lu, and Z.-P. Song. 2007. Phenotypic plasticity rather than locally adapted ecotypes allows the invasive alligator weed to colonize a wide range of habitats. Biological invasions 9: 245–256.

Grimsby, J. L., and R. Kesseli. 2010. Genetic composition of invasive Japanese knotweed s.l. in the United States. Biological invasions 12: 1943–1946.

Grimsby, J. L., D. Tsirelson, M. A. Gammon, and R. Kesseli. 2007. Genetic diversity and clonal vs. sexual reproduction in Fallopia spp. (Polygonaceae). American journal of botany 94: 957–964.

Groeneveld, E., F. Belzile, and C. Lavoie. 2014. Sexual reproduction of Japanese knotweed (Fallopia japonica s.l.) at its northern distribution limit: new evidence of the effect of climate warming on an invasive species. American journal of botany 101: 459–466.

Gross, B. L., and L. H. Rieseberg. 2005. The ecological genetics of homoploid hybrid speciation. The Journal of heredity 96: 241–252.

Hawes, N. A., A. E. Fidler, L. A. Tremblay, X. Pochon, B. J. Dunphy, and K. F. Smith. 2018. Understanding the role of DNA methylation in successful biological invasions: a review. Biological invasions 20: 2285–2300.

Herman, J. J., and S. E. Sultan. 2016. DNA methylation mediates genetic variation for adaptive transgenerational plasticity. Proceedings of the Royal Society B: Biological Sciences 283.

Hollingsworth, M. L., and J. P. Bailey. 2000. Evidence for massive clonal growth in the invasive weed Fallopia japonica (Japanese Knotweed). Botanical journal of the Linnean Society. Linnean Society of London 133: 463–472.

Inamura, A., Y. Ohashi, E. Sato, Y. Yoda, T. Masuzawa, M. Ito, and K. Yoshinaga. 2000. Intraspecific Sequence Variation of Chloroplast DNA Reflecting Variety and Geographical Distribution of Polygonum cuspidatum (Polygonaceae) in Japan. Journal of plant research 113: 419–426.

Jablonka, E., and G. Raz. 2009. Transgenerational epigenetic inheritance: prevalence, mechanisms, and implications for the study of heredity and evolution. The Quarterly review of biology 84: 131–176.

Johnson, P. C. 2014. Extension of Nakagawa & Schielzeth’s R2GLMM to random slopes models. Methods in ecology and evolution /British Ecological Society 5: 944–946.

Johnston, J. A., L. A. Donovan, and M. L. Arnold. 2004. Novel phenotypes among early generation hybrids of two Louisiana iris species: flooding experiments. The Journal of ecology 92: 967–976.

Karrenberg, S., C. Edelist, C. Lexer, and L. Rieseberg. 2006. Response to salinity in the homoploid hybrid species Helianthus paradoxus and its progenitors H. annuus and H. petiolaris. The New phytologist 170: 615–629.

Kawecki, T. J., and D. Ebert. 2004. Conceptual issues in local adaptation. Ecology letters 7: 1225–1241.

Keller, S. R., and D. R. Taylor. 2008. History, chance and adaptation during biological invasion: separating stochastic phenotypic evolution from response to selection. Ecology letters 11: 852–866.

Kingsolver, J. G., H. E. Hoekstra, D. Berrigan, S. N. Vignieri, C. E. Hill, A. Hoang, P. Gibert, and P. Beerli. 2001. The Strength of Phenotypic Selection in Natural Populations. The American Naturalist 157: 245–261.

van Kleunen, M., O. Bossdorf, and W. Dawson. 2018. The Ecology and Evolution of Alien Plants. Annual review of ecology, evolution, and systematics 49: 25–47.

van Kleunen, M., and M. Fischer. 2005. Constraints on the evolution of adaptive phenotypic plasticity in plants. The New phytologist 166: 49–60.

Korner-Nievergelt, F., T. Roth, S. von Felten, J. Guélat, B. Almasi, and P. Korner-Nievergelt. 2015. Bayesian Data Analysis in Ecology Using Linear Models with R, BUGS, and Stan. Academic Press.

Krebs, C., G. Mahy, D. Matthies, U. Schaffner, M.-S. Tiébré, and J.-P. Bizoux. 2010. Taxa distribution and RAPD markers indicate different origin and regional differentiation of hybrids in the invasive Fallopia complex in central-western Europe. Plant biology 12: 215–223.

de Kroon, H., and J.-M. Groenendael. 1997. The Ecology and Evolution of Clonal Plants. Backhuys Publishers.

Lande, R., and S. J. Arnold. 1983. The measurement of selection on correlated characters. Evolution; international journal of organic evolution 37: 1210–1226.

Lavergne, S., and J. Molofsky. 2007. Increased genetic variation and evolutionary potential drive the success of an invasive grass. Proceedings of the National Academy of Sciences of the United States of America 104: 3883–3888.

Lee, C. E. 2002. Evolutionary genetics of invasive species. Trends in ecology & evolution 17: 386–391.

Leger, E. A., and K. J. Rice. 2007. Assessing the speed and predictability of local adaptation in invasive California poppies (Eschscholzia californica). Journal of evolutionary biology 20: 1090–1103.

Leimu, R., and M. Fischer. 2008. A meta-analysis of local adaptation in plants. PloS one 3: e4010.

Lexer, C., M. E. Welch, O. Raymond, and L. H. Rieseberg. 2003. The origin of ecological divergence in Helianthus paradoxus (Asteraceae): selection on transgressive characters in a novel hybrid habitat. Evolution; international journal of organic evolution 57: 1989–2000.

Linhart, Y. B., and M. C. Grant. 1996. Evolutionary significance of local genetic differentiation in plants. Annual review of ecology and systematics 27: 237–277.

Liu, W., Y. Zhang, X. Chen, K. Maung-Douglass, D. R. Strong, and S. C. Pennings. 2020. Contrasting plant adaptation strategies to latitude in the native and invasive range of Spartina alterniflora. The New phytologist 226: 623–634.

Long, R. 2021. Crossed random effects: how do we model multiple reciprocal transplants in lme4? *stackexchange.com*. Website https://stats.stackexchange.com/users/7486/robert-long), Crossed random effects: how do we model multiple reciprocal transplants in lme4?, URL (version: 2021-06-16): https://stats.stackexchange.com/q/530923 [accessed 17 June 2021].

Loomis, E. S., and L. Fishman. 2009. A Continent-Wide Clone: Population Genetic Variation of the Invasive Plant Hieracium aurantiacum (Orange Hawkweed; Asteraceae) in North America. International journal of plant sciences 170: 759–765.

Mandák, B., K. Bímová, P. Pyšek, J. Štěpánek, and I. Plačková. 2005. Isoenzyme diversity in Reynoutria (Polygonaceae) taxa: escape from sterility by hybridization. Plant systematics and evolution = Entwicklungsgeschichte und Systematikder Pflanzen 253: 219–230.

Mandák, B., P. Pyšek, K. Bímová, and Others. 2004. History of the invasion and distribution of Reynoutria taxa in the Czech Republic: a hybrid spreading faster than its parents. Preslia 76: 15–64.

Matesanz, S., T. Horgan-Kobelski, and S. E. Sultan. 2015. Evidence for rapid ecological range expansion in a newly invasive plant. AoB plants 7.

Matesanz, S., and S. E. Sultan. 2013. High-performance genotypes in an introduced plant: insights to future invasiveness. Ecology 94: 2464–2474.

Mauricio, R., and M. D. Rausher. 1997. Experimental manipulation of putative selective agents provides evidence for the role of natural enemies in the evolution of plant defense. Evolution; international journal of organic evolution 51: 1435–1444.

Mounger, J., M. L. Ainouche, O. Bossdorf, A. Cavé-Radet, B. Li, M. Parepa, A. Salmon, et al. 2021. Epigenetics and the success of invasive plants. Philosophical transactions of the Royal Society of London. Series B, Biological sciences 376: 20200117.

Murren, C. J., J. R. Auld, H. Callahan, C. K. Ghalambor, C. A. Handelsman, M. A. Heskel, J. G. Kingsolver, et al. 2015. Constraints on the evolution of phenotypic plasticity: limits and costs of phenotype and plasticity. Heredity 115: 293–301.

Muth, N. Z., and M. Pigliucci. 2007. Implementation of a novel framework for assessing species plasticity in biological invasions: responses of Centaurea and Crepis to phosphorus and water availability. The Journal of ecology 95: 1001–1013.

Nakagawa, S., P. C. D. Johnson, and H. Schielzeth. 2017. The coefficient of determination R2 and intra-class correlation coefficient from generalized linear mixed-effects models revisited and expanded. Journal of the Royal Society, Interface / the Royal Society 14.

Nakagawa, S., and H. Schielzeth. 2013. A general and simple method for obtainingR2from generalized linear mixed-effects models. Methods in ecology and evolution /British Ecological Society 4: 133–142.

Neinavaie, F., A. Ibrahim-Hashim, A.M. Kramer, J.S. Brown, and C.L. Richards. 2021. The genomic processes of biological invasions: from invasive species to cancer metastases and back again. Frontiers in Ecology and Evolution 9: Article 681100.

Nimon, K., M. Lewis, R. Kane, and R. M. Haynes. 2008. An R package to compute commonality coefficients in the multiple regression case: an introduction to the package and a practical example. Behavior research methods 40: 457–466.

Oduor, A. M. O., R. Leimu, and M. van Kleunen. 2016. Invasive plant species are locally adapted just as frequently and at least as strongly as native plant species. The Journal of ecology 104: 957–968.

Oplaat, C., and K. J. F. Verhoeven. 2015. Range expansion in asexual dandelions: selection for general-purpose genotypes? The Journal of ecology 103: 261–268.

Pan, X., Y. Geng, W. Zhang, B. Li, and J. Chen. 2006. The influence of abiotic stress and phenotypic plasticity on the distribution of invasive Alternanthera philoxeroides along a riparian zone. Acta Oecologica 30: 333–341.

Papadopulos, A. S. T., A. J. Helmstetter, O. G. Osborne, A. A. Comeault, D. P. Wood, E. A. Straw, L. Mason, et al. 2021. Rapid Parallel Adaptation to Anthropogenic Heavy Metal Pollution. Molecular biology and evolution.

Parepa, M., M. Fischer, C. Krebs, and O. Bossdorf. 2014. Hybridization increases invasive knotweed success. Evolutionary applications 7: 413–420.

Park, C.-W., G. S. Bhandari, H. Won, J. H. Park, and D. S. Park. 2018. Polyploidy and introgression in invasive giant knotweed (Fallopia sachalinensis) during the colonization of remote volcanic islands. Scientific reports 8: 16021.

Prentis, P. J., J. R. U. Wilson, E. E. Dormontt, D. M. Richardson, and A. J. Lowe. 2008. Adaptive evolution in invasive species. Trends in plant science 13: 288–294.

Puy, J., F. de Bello, H. Dvořáková, N. G. Medina, V. Latzel, and C. P. Carmona. 2021. Competition-induced transgenerational plasticity influences competitive interactions and leaf decomposition of offspring. The new phytologist 229: 3497–3507.

Puy, J., C. P. Carmona, H. Dvořáková, V. Latzel, and F. de Bello. 2021. Diversity of parental environments increases phenotypic variation in Arabidopsis populations more than genetic diversity but similarly affects productivity. Annals of botany 127: 425–436.

Pysek, P., J. H. Brock, K. Bímová, B. Mandák, V. Jarosík, I. Koukolíková, J. Pergl, and J. Stepánek. 2003. Vegetative regeneration in invasive Reynoutria (Polygonaceae) taxa: the determinant of invasibility at the genotype level. American journal of botany 90: 1487–1495.

Pyšek, P., B. Mandák, T. Francírková, and K. Prach. 2001. Persistence of stout clonal herbs as invaders in the landscape: A field test of historical records. In G. Brundu, J. Brock, I. Camarda, L. Child, and M. Wade [eds.], Plant Invasions: Species Ecology and Ecosystem Management., 235–244. Backhuys Publishers.

Qiao, H., W. Liu, Y. Zhang, Y.-Y. Zhang, and Q. Q. Li. 2019. Genetic admixture accelerates invasion via provisioning rapid adaptive evolution. Molecular ecology 28: 4012–4027.

Rapp, R. A., and J. F. Wendel. 2005. Epigenetics and plant evolution. The New phytologist 168: 81–91.

R Core Team. 2020. R: A language and environment for statistical computing.

Reimann, C., and S.-W. Breckle. 1995. Salt tolerance and ion relations of Salsola kali L.: differences between ssp. tragus (L.) Nyman and ssp. ruthenica (Iljin) Soó. The New phytologist 130: 37–45.

Richards, C. L., C. Alonso, C. Becker, O. Bossdorf, E. Bucher, M. Colomé-Tatché, W. Durka, et al. 2017. Ecological plant epigenetics: Evidence from model and non-model species, and the way forward. Ecology letters 20: 1576–1590.

Richards, C. L., O. Bossdorf, N. Z. Muth, J. Gurevitch, and M. Pigliucci. 2006. Jack of all trades, master of some? On the role of phenotypic plasticity in plant invasions. Ecology letters 9: 981–993.

Richards, C. L., and M. Pigliucci. 2020. Epigenetic Inheritance. A Decade into the Extended Evolutionary Synthesis. Paradigmi 38: 463–494.

Richards, C. L., A. W. Schrey, and M. Pigliucci. 2012. Invasion of diverse habitats by few Japanese knotweed genotypes is correlated with epigenetic differentiation. Ecology letters 15: 1016–1025.

Richards, C. L., R. L. Walls, and J. P. Bailey. 2008. Plasticity in salt tolerance traits allows for invasion of novel habitat by Japanese knotweed sl (Fallopia japonica and F.×bohemica, Polygonaceae). American Journal of.

Richards, C. L., S. N. White, M. A. McGuire, and S. J. Franks. 2010. Plasticity, not adaptation to salt level, explains variation along a salinity gradient in a salt marsh perennial. Estuaries.

Roff, D. A., and T. Mousseau. 2005. The evolution of the phenotypic covariance matrix: evidence for selection and drift in Melanoplus. Journal of evolutionary biology 18: 1104–1114.

Rosenthal, D. M., A. P. Ramakrishnan, and M. B. Cruzan. 2008. Evidence for multiple sources of invasion and intraspecific hybridization in Brachypodium sylvaticum (Hudson) Beauv. in North America. Molecular ecology 17: 4657–4669.

Rosenthal, D. M., A. E. Schwarzbach, L. A. Donovan, O. Raymond, and L. H. Rieseberg. 2002. Phenotypic Differentiation between Three Ancient Hybrid Taxa and Their Parental Species. International journal of plant sciences 163: 387–398.

Sakai, A. K., F. W. Allendorf, and J. S. Holt. 2001. The population biology of invasive species. Annual review of.

Salmon, A., M. L. Ainouche, and J. F. Wendel. 2005. Genetic and epigenetic consequences of recent hybridization and polyploidy in Spartina (Poaceae). Molecular ecology 14: 1163–1175.

Sambatti, J. B. M., and K. J. Rice. 2006. Local adaptation, patterns of selection, and gene flow in the Californian serpentine sunflower (Helianthus exilis). Evolution; international journal of organic evolution 60: 696–710.

Schuster, T. M., J. L. Reveal, and K. A. Kron. 2011. Phylogeny of Polygoneae (Polygonaceae: Polygonoideae). Taxon 60: 1653–1666.

Schuster, T. M., K. L. Wilson, and K. A. Kron. 2011. Phylogenetic Relationships of Muehlenbeckia, Fallopia, and Reynoutria (Polygonaceae) Investigated with Chloroplast and Nuclear Sequence Data. International journal of plant sciences 172: 1053–1066.

Sexton, J. P., J. K. McKay, and A. Sala. 2002. Plasticity and genetic diversity may allow saltcedar to invade cold climates in north America. Ecological applications: a publication of the Ecological Society of America 12: 1652.

Shi, W., X. Chen, L. Gao, C.-Y. Xu, X. Ou, O. Bossdorf, J. Yang, and Y. Geng. 2018. Transient Stability of Epigenetic Population Differentiation in a Clonal Invader. Frontiers in plant science 9: 1851.

Stoffel, M. A., S. Nakagawa, and H. Schielzeth. 2017. rptR: repeatability estimation and variance decomposition by generalized linear mixed-effects models. Methods in ecology and evolution / British Ecological Society 8: 1639–1644.

Sultan, S. E. 2004. Promising directions in plant phenotypic plasticity. Perspectives in plant ecology, evolution and systematics 6: 227–233.

Sultan, S. E., T. Horgan-Kobelski, L. M. Nichols, C. E. Riggs, and R. K. Waples. 2013. A resurrection study reveals rapid adaptive evolution within populations of an invasive plant. Evolutionary applications 6: 266–278.

Tiébré, M.-S., J.-P. Bizoux, O. J. Hardy, J. P. Bailey, and G. Mahy. 2007. Hybridization and morphogenetic variation in the invasive alien Fallopia (Polygonaceae) complex in Belgium. American journal of botany 94: 1900–1910.

Townsend, A. 1997. Japanese Knotweed: A Reputation Lost. Arnoldia Zimbabwe 57: 13–19.

Van Kleunen, M., and M. Fischer. 2007. Progress in the detection of costs of phenotypic plasticity in plants. The New phytologist 176: 727–730.

VanWallendael, A., M. Alvarez, and S. J. Franks. 2021. Patterns of population genomic diversity in the invasive Japanese knotweed species complex. American journal of botany 108: 857–868.

VanWallendael, A., E. Hamann, and S. J. Franks. 2018. Evidence for plasticity, but not local adaptation, in invasive Japanese knotweed (Reynoutria japonica) in North America. Evolutionary ecology 32: 395–410.

Verhoeven, K. J. F., J. J. Jansen, P. J. Van Dijk, and A. Biere. 2010. Stress-induced DNA methylation changes and their heritability in asexual dandelions. The New phytologist 185: 1108–1118.

Walls, R. L. 2010. Hybridization and Plasticity Contribute to Divergence Among Coastal and Wetland Populations of Invasive Hybrid Japanese Knotweed s.l. (Fallopia spp.). Estuaries and Coasts 33: 902–918.

Weber, E., and C. M. D’Antonio. 1999. Germination and growth responses of hybridizing Carpobrotus species (Aizoaceae) from coastal California to soil salinity. American journal of botany 86: 1257–1263.

Zerebecki, R. A., E. E. Sotka, T. C. Hanley, K. L. Bell, C. Gehring, C. C. Nice, C. L. Richards, and A. R. Hughes. 2021. Repeated Genetic and Adaptive Phenotypic Divergence across Tidal Elevation in a Foundation Plant Species. The American naturalist: E000–E000.

Zhang, Y.-Y., M. Parepa, M. Fischer, and O. Bossdorf. 2016. Epigenetics of colonizing species? A study of Japanese knotweed in Central Europe. In S. Barrett, R. I. Colautti, K. M. Dlugosch, and L. H. Rieseberg [eds.], Invasion Genetics: The Baker and Stebbins Legacy, 328–340. Wiley-Blackwell UK.

